# FGFR inhibitor mediated dismissal of SWI/SNF complexes from YAP-dependent enhancers induces adaptive therapeutic resistance

**DOI:** 10.1101/2021.03.02.433446

**Authors:** Yihao Li, Xintao Qiu, Xiaoqing Wang, Hui Liu, Renee C. Geck, Alok K. Tewari, Tengfei Xiao, Alba Font-Tello, Klothilda Lim, Kristen L. Jones, Murry Morrow, Raga Vadhi, Pei-Lun Kao, Aliya Jaber, Smitha Yerrum, Yingtian Xie, Kin-Hoe Chow, Paloma Cejas, Quang-Dé Nguyen, Henry W. Long, X. Shirley Liu, Alex Toker, Myles Brown

**Affiliations:** Department of Medical Oncology, Dana-Farber Cancer Institute, Harvard Medical School, Boston, MA 02115, USA; Center for Functional Cancer Epigenetics, Dana-Farber Cancer Institute, Boston, MA 02115, USA; Department of Pathology, and Cancer Center, Beth Israel Deaconess Medical Center, Harvard Medical School, Boston, MA 02115, USA; Department of Oncologic Pathology, Dana-Farber Cancer Institute, Boston, MA 02115, USA; Center for Patient Derived Models, Dana-Farber Cancer Institute, Boston, MA 02115, USA; Lurie Family Imaging Center, Center for Biomedical Imaging in Oncology, Dana-Farber Cancer Institute, Boston, Massachusetts 02210, USA; Department of Biostatistics and Computational Biology, Dana-Farber Cancer Institute, Harvard T.H. Chan School of Public Health, Boston, MA 02115, USA; Ludwig Center at Harvard, Harvard Medical School, Boston, MA 02215, USA

## Abstract

How cancer cells adapt to evade the therapeutic effects of drugs targeting oncogenic drivers is poorly understood. Here we report an epigenetic mechanism leading to the adaptive resistance of triple-negative breast cancer (TNBC) to fibroblast growth factor receptor (FGFR) inhibitors. Prolonged FGFR inhibition suppresses the function of BRG1-dependent chromatin remodeling leading to an epigenetic state that derepresses YAP-associated enhancers. These chromatin changes induce the expression of several amino acid transporters resulting in increased intracellular levels of specific amino acids that reactivate mTORC1. Collectively, these findings reveal a novel feedback loop involving an epigenetic state transition and metabolic reprogramming that leads to adaptive therapeutic resistance.

## Introduction

Improving therapeutic options for triple negative breast cancer (TNBC) is a major unmet need in breast cancer treatment. TNBC are a heterogeneous breast cancer subtype characterized by the lack of expression of estrogen receptor (ER), progesterone receptor (PR) or amplification of human epidermal growth factor 2 receptor (HER2/ERBB2) and thus far, few specific targeted therapies have been shown to achieve durable responses in TNBC patients. While there are few recurrent somatic genetic alterations in this breast cancer subtype, fibroblast growth factor receptors 1-3 (FGFR 1-3) have been shown to be amplified in approximately 10-15% ^1-3^ of cases. FGFR1 and 2 alterations are associated with increased risk of distant metastasis and worse overall survival in breast cancer and have been suggested to be therapeutic targets ^3-5^. Aberrant activation of FGF receptors plays a central role in tumor growth, survival, epithelial-mesenchymal transition (EMT), angiogenesis, and metastasis in various solid tumors ^6^. FGF signaling is stimulated by FGF ligands binding to their cognate receptors, and subsequent receptor phosphorylation activates downstream growth and survival pathways including RAS-MAPK, PI3K-AKT and JAK-STAT ^7^. Pre-clinical studies have demonstrated that treatment of TNBC cells with FGFR inhibitors *in vitro* and *in vivo* effectively blocks tumor cell proliferation relative to other subtypes of breast cancer ^8, 9^. Previous work has identified FGF signaling as oncogenic drivers of mouse TNBC, and these tumors are sensitive to FGFR inhibition ^10^.

Therapies targeting FGFRs have shown promising results in many cancer types and been approved for the treatment of urothelial carcinoma and cholangiocarcinoma ^11, 12^. Several FGFR inhibitors have been evaluated in clinical trials across multiple solid tumors including triple negative breast cancer (NCT04024436) ^13-15^. However, adaptive or intrinsic resistance is a common problem limiting therapeutic efficacy though the mechanisms leading to resistance are poorly understood. Here, we integrated genetic screens, bulk and single cell epigenomic, proteomic and metabolomic profiling to define the mechanisms mediating adaptive resistance to FGFR inhibitors and identify potential effective therapeutic strategies for treating FGFR-aberrant TNBCs.

## Results

### Identification of sensitizers to FGFR inhibitors in TNBC cells by genome-wide CRISPR screens

As previous clinical analyses have demonstrated FGFR altercations in TNBCs ^1-3^, we tested the efficacy and specificity of infigratinib, a potent and selective FGFR inhibitor currently applied in clinical trials, using multiple breast cancer cell lines with or without aberrant FGFR1/2 expression (Extended Data Fig. 1a) ^16^. Infigratinib specifically decreased FGFR-aberrant TNBC cell proliferation (MEM223, CAL-120, Hs578T). In contrast, other non-FGFR altered cells were resistant to infigratinib treatment (Fig. 1a).

**Fig. 1.**
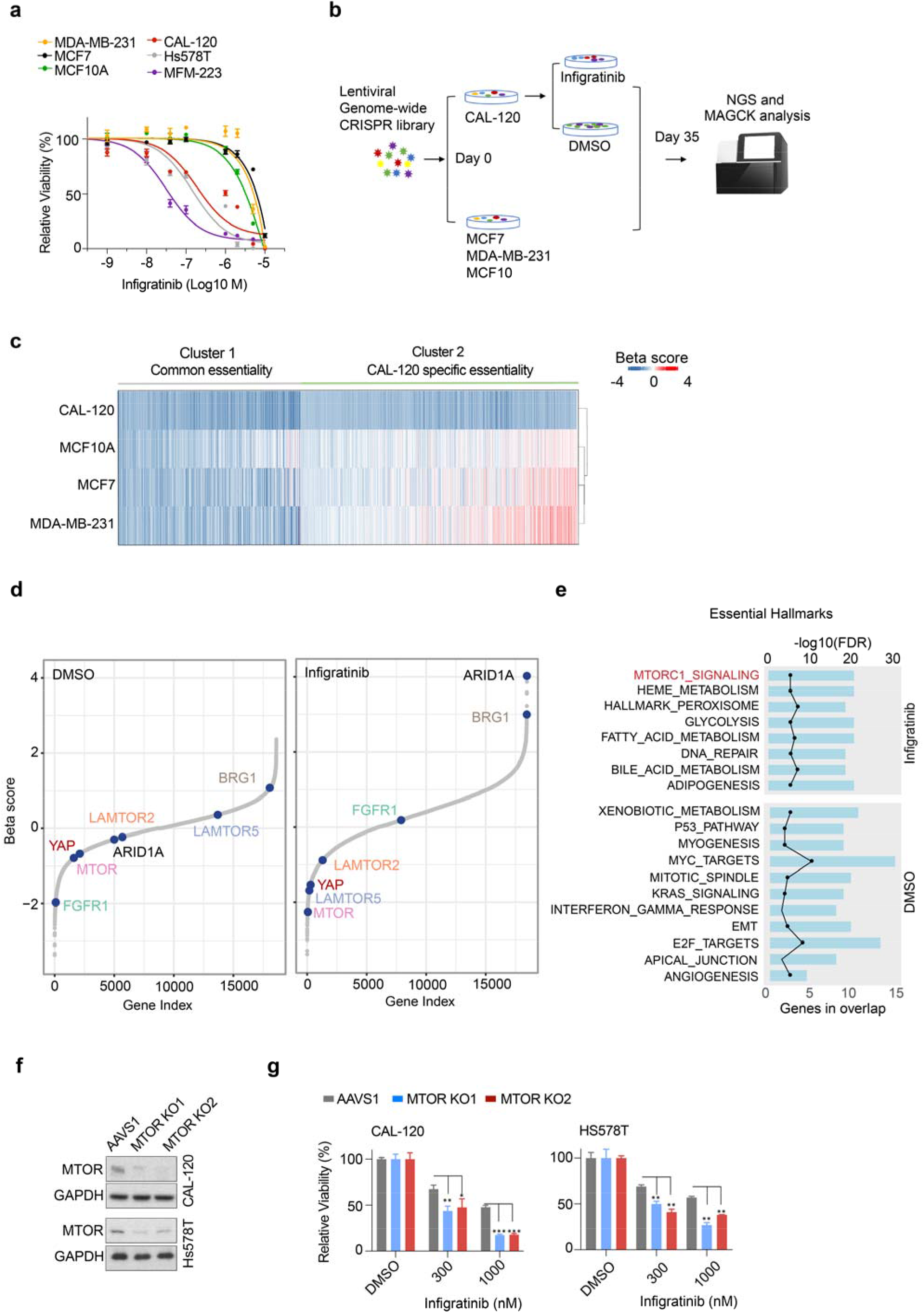
Identification of vulnerabilities of FGFR-aberrant TNBC cells. **a**, Dose response of MDA-MB-231, MCF7, MCF10A, CAL-120, Hs578T and MFM-223 cells treated with infigratinib for 6 days. **b**, Schematic of the genome-wide CRISPR knock out screen in the indicated cell lines. **c**, Heatmap of the genetic dependencies for the top 1500 essential genes in CAL-120 cells across multiple cell lines. Genes are clustered according to common essential genes (genes that are essential in more than one cell line) and CAL-120 specific essential genes (genes that are only essential in the CAL-120 cell line). The negative beta scores represent genes whose sgRNA are depleted upon completion of the screen, and the positive beta scores represent genes whose sgRNA are enriched upon completion of the screen. **d**, Rank of gene essentiality of genome-wide CRISPR screen in CAL-120 cells in the presence of DMSO (left) or 300 nM infigratinib (right). The key dependencies in each condition are marked as indicated. **e**, Plot of significantly enriched hallmark gene sets based on essential genes in CAL-120 cells treated with infigratinib versus DMSO. **f**, Immunoblot of mTOR and GAPDH protein levels in CAL-120 and Hs578T cells harboring CRISPR sgRNA mediated knockout of mTOR or AAVS1 control. **g**, The relative viability of CAL-120 and Hs578T cells harboring CRISPR knock out of AAVS1 (control) or mTOR in presence of DMSO, 300 nM, or 1000 nM infigratinib for 6 days. Data presented as mean± SD. \**p*<0.05, \*\**p*<0.01, \*\*\**p*<0.001.

Since CAL-120 cells displayed an intermediate response to FGFR inhibition, we utilized CRISPR knock out (KO) screens in this model treated with or without infigratinib to detect potential mechanisms of resistance to FGFR inhibitors. In parallel, CRISPR KO screens were also performed in MDA-MB-231, MCF7, and MCF10A cells to allow identification of genetic dependencies unique to FGFR-driven TNBCs (Fig. 1b). In our system, the human whole-genome CRISPR guide library targets more than 18,000 genes, with 6 guide RNAs per gene ^17^. After infection with the lentiviral library, selection with puromycin, and exposure to vehicle or drug, cells were collected at day 0 and day 35 for deep sequencing (Extended Data Fig. 1b). Thereafter, we analyzed sgRNA abundance and quantified gene essentiality by MAGeCK-VISPR ^18^, with essential genes having a more negative beta score. To further refine our candidate list of essential genes in FGFR-driven TNBC cells, we performed comparison of the top essential genes from CAL-120 cells in all cell lines (Fig. 1c and Supplementary Table 1). We found that about 40% of CAL-120 essential genes (cluster 1) are common amongst the breast cancer cell lines tested, whereas the remaining genes (cluster 2) contained the essential genes unique to CAL-120 cells. The common essential genes in cluster 1 were enriched for RNA/DNA processing and cell cycle functions (Extended Data Fig. 1c). As expected, FGFR pathway signatures were significantly enriched in the set of CAL-120 essential genes (Extended Data Fig. 1d).

To identify those genes that become more essential in the presence of FGFR inhibition, we analyzed genetic dependencies in the CAL-120 cells in the absence or presence of 300 nM infigratinib, a dose causes approximately a 30% inhibition of cell growth. As shown in Figure 1d, FGFR1 is one of the top essential genes in the CAL-120 cells grown under vehicle conditions (left panel). In contrast, in the presence of infigratinib, FGFR1 becomes non-essential with a beta score near 0 (right panel), supporting the FGFR-selective action of infigratinib. Interestingly, components of the mTORC1 signal pathway including mTOR, LAMTOR2 and LAMTOR5 become more essential upon infigratinib treatment. In addition, a key component of Hippo signaling, YAP, was also more essential in CAL-120 cells upon FGFR inhibition. In contrast, the SNI/SNF chromatin remodeling complex factors ARID1A and BRG1 (SMARCA4) genes are positively selected in the presence of the FGFR inhibitor suggesting that their loss contributes to resistance to this drug. The genes that were more essential in the presence of the FGFR inhibitor were significantly enriched in mTORC1 signaling, whereas the essential genes for growth in DMSO treated cells were enriched for E2F, MYC and the KRAS signaling (Fig. 1e). We therefore tested whether mTORC1 signal repression could sensitize FGFR-driven cells to FGFR inhibition by depleting mTOR levels in CAL-120, Hs578T and MFM-223 cells (Fig. 1f and Extended Data Fig. 1e). Consistent with our CRISPR screen results, mTOR depletion significantly impaired cell proliferation upon infigratinib treatment (Fig. 1g and Extended Data Fig. 1f). Together, these results identify the mTORC1 pathway as a genetic dependency in TNBC cells upon FGFR inhibition.

### AKT/MAPK independent feedback activation of mTORC1 determines adaptive resistance to FGFR inhibition

To explore the process of adaptive resistance to infigratinib, we treated CAL-120, Hs578T and MFM223 cells with infigratinib for various times spanning from one to twelve days and analyzed FGFR and its downstream signaling at each time point. Infigratinib disrupted FGFR phosphorylation in all cell lines at all time points. While AKT phosphorylation was unchanged by infigratinib treatment in the CAL-120 and Hs578T cells driven by FGFR1 (Fig. 2a), it was inhibited in MFM223 cells driven by FGFR2 as previously reported (Extended Data Fig. 2a) ^19^. Short-term FGFR inhibition for one or three days efficiently decreased phosphorylation of ERK and the mTORC1 downstream targets S6K and S6 in all cell lines, ERK, S6K and S6 phosphorylation reappeared upon prolonged FGFR inhibition in all cell lines tested (Fig. 2a and Extended Data Fig. 2a). Interestingly, this event occurred in the absence of AKT phosphorylation even in the FGFR2-driven MFM223 cells, suggesting that mTOR reactivation is independent of AKT. Treatment with dovitinib, a distinct

**Fig. 2.**
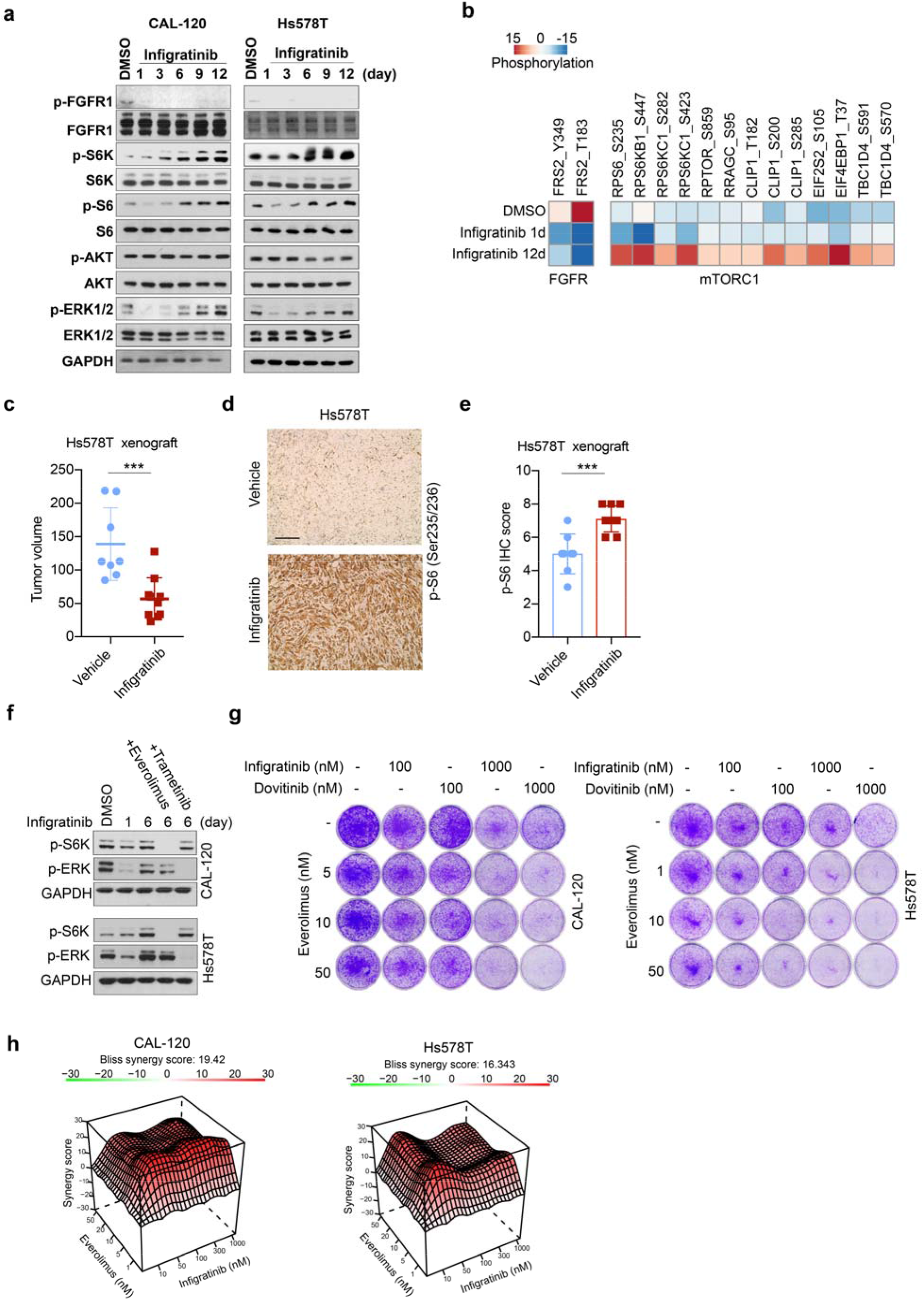
Reactivation of mTORC1 signaling is required for adaptive resistance to FGFR inhibitors. **a**, Immunoblot of FGFR1, ERK1/2, AKT, S6K, and S6 protein phosphorylation and total levels in CAL-120 and Hs578T cells in the presence of infigratinib (300 nM) at indicated timepoints. **b**, Phosphoproteomic analysis of protein phosphorylation in CAL-120 cells treated with DMSO or infigratinib for 1 or 12 days. Heatmap depicting relative phosphorylation levels of mTORC1 and FGFR substrates at indicated sites. **c**, Tumor volumes of Hs578T mouse xenografts treated with vehicle or infigratinib for 35 days. \*\*\**p*<0.001. **d**, Immunohistochemical (IHC) staining of pS6 (Ser235/236) in Hs578T tumors treated with vehicle or infigratinib (12.5 mg/kg) for 35 days. Scale bar: 100 μm. **e**, Quantification of pS6 (Ser235/236) IHC scores in Hs578T tumors treated with or without infigratinib for 35 days. **f**, Immunoblot of ERK 1/2 and S6K phosphoprotein levels in CAL-120 (upper) and Hs578T (lower) cells treated for 6d with infigratinib (300 nM), alone and in combination with 50 nM everolimus or 50 nM trametinib for 24 hours. **g**, Clonogenic assay of CAL-120 and Hs578T cells treated with infigratinib, dovitinib, everolimus alone or in the combinations indicated for 12 days. **h**, Bliss model of infigratinib and everolimus synergy in CAL-120 and Hs578T cells. Positive score represents a synergistic effect.

FGFR inhibitor, showed similar results (Extended Data Fig. 2b). To evaluate whether alterations in protein phosphorylation are relevant for feedback activation of mTORC1, we performed phospho-proteomic profiling on CAL-120 cells either untreated or treated with infigratinib for 1 or 12 days (Supplementary Table 2). We found that the phosphorylation levels of many known mTORC1 substrates were relatively low under control conditions and following short-term infigratinib treatment while these phospho-proteins were dramatically upregulated following long-term FGFR inhibition (Fig. 2b) ^20^.

Next, we examined the ability of infigratinib to increase mTORC1 signaling *in vivo*. In Hs578T xenografts, infigratinib treatment significantly delayed tumor growth (Fig. 2c), however, S6 phosphorylation was dramatically increased at the same time point (Fig. 2d, e).

Considering that activation of ERK signaling also signals through mTORC1 ^21^ and increased phosphorylation of ERK is observed with long-term drug FGFR inhibitor treatment, we tested the crosstalk between ERK, mTORC1 and FGFR pharmacologically in CAL-120 and Hs578T cell lines (Fig. 2f). The mTORC1 inhibitor everolimus efficiently blocked infigratinib-mediated feedback phosphorylation of S6K, but had no effect on ERK phosphorylation. Although the MEK inhibitor trametinib impeded infigratinib-induced ERK phosphorylation, it did not interfere with S6K phosphorylation. This result demonstrates that the feedback loop involving mTORC1 is ERK independent. Thus, we next tested whether mTOR and FGFR inhibitors together could more effectively block cell proliferation in FGFR-driven TNBCs. CAL-120 and Hs578T cells were treated with either FGFR inhibitors (infigratinib or dovitinib) or mTOR inhibitor (everolimus) alone, or in combination, and the impact on cell colony formation was assessed. Though FGFR inhibition alone decreased cell proliferation, combined treatment with everolimus lead to dramatic growth inhibition in both cell lines (Fig. 2g). Moreover, combining infigratinib and everolimus showed significant synergy as assessed by Bliss synergy analysis in CAL-120, Hs578T, and MFM223 cells (Fig. 2h and Extended Data Fig. 2c). The third generation mTOR inhibitor Rapalink-1 also demonstrated strong synergy with infigratinib in CAL-120 cells (Extended Data Fig. 2d). These results demonstrate that treatment of FGFR-driven TNBC with FGFR inhibitors activate a feedback loop leading to ERK- and AKT-independent mTORC1 activation that increases cell growth and survival, thereby limiting FGFR inhibitor efficacy.

### FGFR inhibition increases cellular levels of amino acids

To identify the regulatory mechanisms that contribute to feedback activation of mTORC1 upon FGFR inhibition, we examined changes in gene expression over a time course of infigratinib treatment in CAL-120 cells (Supplementary Table 3). We observed significant differences in gene expression with drug treatment. While some genes were down regulated across all time points, the expression pattern of a substantial number of genes that were either induced or repressed upon 4h FGFR inhibition began to switch at 48h and were dramatically reversed following 12d of FGFR inhibition (Fig. 3a). We then analyzed the impact of prolonged infigratinib treatment on global protein levels by proteomic profiling in CAL-120 cells (Supplementary Table 2). In agreement with our RNA-seq data, 1 day of drug treatment altered protein expression patterns in CAL-120 cells, with a number of proteins reversing expression patterns after 12 days of treatment (Extended Data Fig. 3a). We observed a strong and significant correlation between long-term changes of mRNA and protein levels in response to infigratinib (Fig. 3b), and interestingly identified both YAP target genes and several amino acid transporters as some of the most dramatically upregulated transcripts and proteins ^22-24^. This finding was reinforced by gene set enrichment analysis (GSEA) of both RNA and protein data from CAL-120 cells upon long-term FGFR inhibition (Fig. 3c and Extended Data Fig. 3b).

**Fig. 3.**
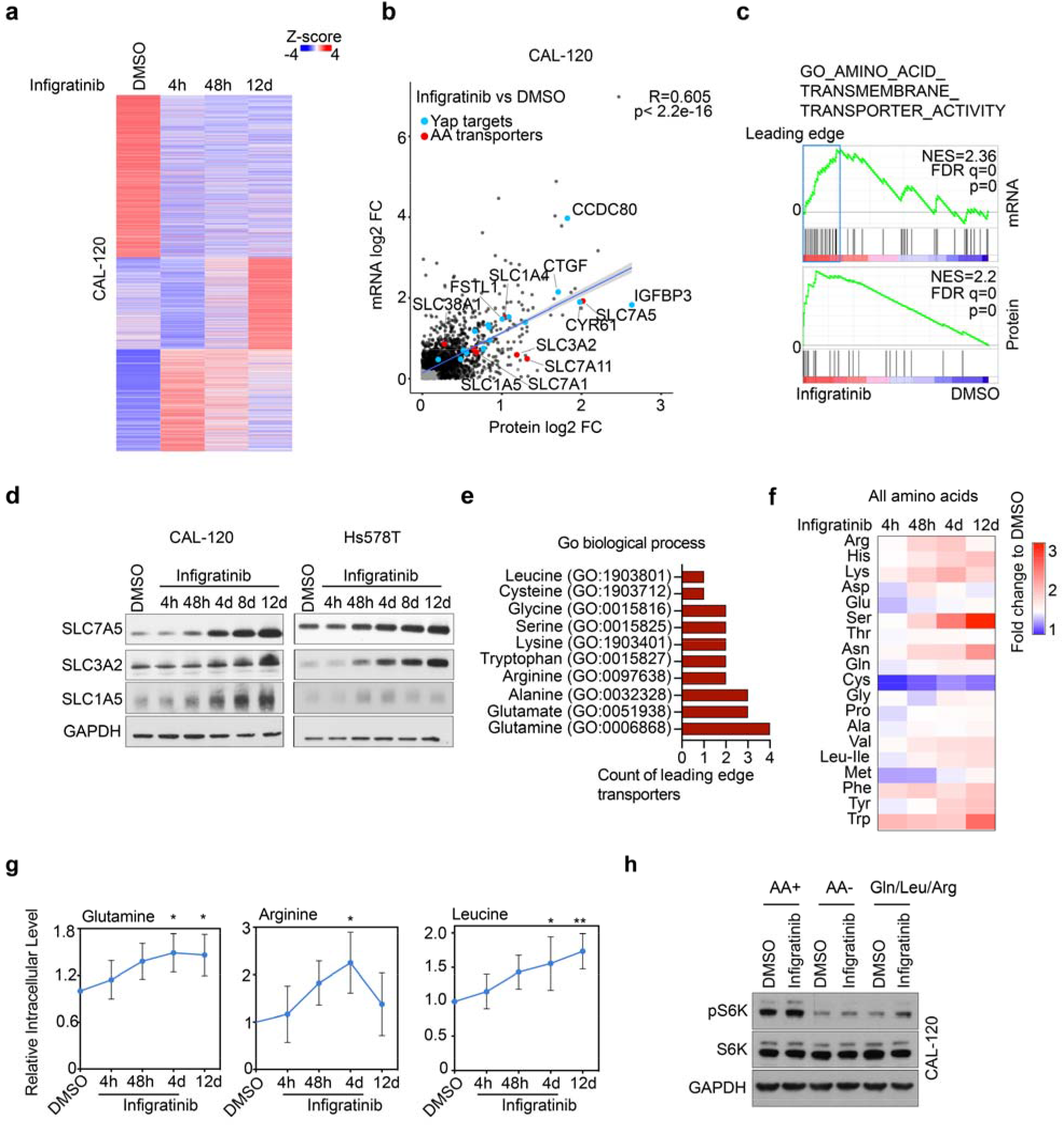
Upregulation of amino acid transporters drives mTORC1 signaling upon FGFR inhibition. **a**, K-means clustering of the most differentially regulated genes (n=5548) in CAL-120 cells treated with infigratinib versus DMSO. Heatmap displays the gene expression of CAL-120 cells treated with DMSO or infigratinib for 4h, 48h, or 12d. **b**, Scatterplot showing log2 fold changes of significantly increased (black dots) mRNA and proteins in CAL-120 cells treated with 12d infigratinib (300 nM) versus DMSO. Conserved significantly changed YAP target genes and amino acid transporters are marked in blue and red respectively. **c**, GSEA analysis of amino acid transmembrane transporter signature in differentially expressed mRNA (upper) or protein (lower) levels in CAL120 cells treated with 12d infigratinib versus DMSO. **d**, Immunoblot of SLC7A5, SLC3A2, and SLC1A5 in CAL-120 and Hs578T cells in the presence of DMSO or 300 nM infigratinib at indicated time points. **e**, GO term analysis of amino acids that are transported by leading edge amino acid transporters in b. **f**, Heatmap representing fold change of amino acid uptake in CAL-120 cells treated with 4h, 48h, or 12d infigratinib (300 nM) versus DMSO. **g**, Relative cellular levels of glutamine, arginine, and leucine in CAL-120 cells in the presence of infigratinib at indicated time points. \**p*<0.05, \*\**p*<0.01. **h**, Immunoblot of p-S6K and total S6K in CAL-120 cells treated with DMSO or infigratinib (300 nM) for 6 days, cultured in full medium (AA+), amino acid-free medium (AA-), or amino acid-free medium containing glutamine, leucine, and arginine (Gln/Leu/Arg) for 6 hours.

As intracellular amino acids can stimulate mTORC1 activity, we postulated that increased expression of amino acid transporters might be involved in the feedback loop mediating resistance to FGFR inhibition ^25^. We confirmed increased expression of SLC7A5, SLC3A2 and SLC1A5 by immunoblotting (Fig. 3d), which are the amino acid transporters critical for mTORC1 activation ^26^. We found that the amino acid transporters driving the enrichment of the gene set shown in Figure 3c (leading edge) are involved with the transport of amino acids upstream of mTORC1, including glutamine, arginine and leucine ^27-29^ (Fig. 3e). Importantly, these amino acid transporters were upregulated by FGFR inhibition by 2 days of treatment, whereas mTORC1 activation was observed at later points (Fig. 2a), indicating that mTORC1 activation might be a downstream consequence of the increase in amino acid transporter expression.

To determine whether induction of amino acid transporters was associated with upregulated cellular level of amino acids upon FGFR inhibition, we measured cellular metabolites in CAL-120 cells using mass spectrometry (Supplementary Table 4). Several amino acids including glutamine, arginine and leucine, which are implicated in mTORC1 sensing, were increased intracellularly by FGFR inhibition (Fig. 3f, g). To investigate the dependency of mTORC1 feedback activation on these three amino acids, we treated CAL-120 cells in regular media with or without infigratinib for 6 days, and subsequently the cells were cultured for 6 hours with either full media, media depleted of amino acids or depleted media supplemented with glutamine, arginine and leucine. Amino acid depletion inhibited both basal S6K phosphorylation as well as feedback activation of mTORC1 by FGFR inhibition. The addition of glutamine, arginine and leucine to amino acid depleted media partially rescued the feedback activation of mTORC1 as measured by S6K phosphorylation (Fig. 3h). These findings reveal that resistance to FGFR inhibition is mediated, at least in part, by increased amino acid transport resulting in mTORC1 reactivation.

### Enhancer reactivation drives transcriptional changes upon FGFR inhibition

To identify whether the observed mRNA and protein expression changes upon FGFR inhibition are driven by chromatin alterations, we next investigated the landscape of active enhancers by H3K27ac chromatin immunoprecipitation sequencing (ChIP-seq). ChIP-seq was performed in CAL-120 cells either untreated (DMSO) or treated with 300 nM infigratinib for 4h, 48h or 12 days. Long term treatment with infigratinib resulted in 1250 regions of significantly gained H3K27ac signal, whereas only 278 regions were significantly decreased (Extended Data Fig. 4a). Consistent with RNA expression changes, acute FGFR inhibition for 4h reduced the H3K27ac signal, whereas the reactivation of a significant number of enhancers was observed at 48h, and was even more dramatically increased after 12 days of treatment (Fig. 4a). Gained enhancers were correlated with increased transcription of nearby genes, whereas loss of H3K27ac signal was associated with downregulated expression of nearby genes (Fig. 4b). Using the Cistrome Data Browser (CistromeDB) toolkit ^30^, we investigated the similarity between H3K27ac bound regions induced by infigratinib treatment and publicly available transcription factor and chromatin regulator binding sites defined by ChIP-seq. Enhancers with gained H3K27ac signal upon FGFR inhibitor treatment were significantly similar to the DNA binding profiles of Hippo pathway transcriptional components including YAP1, TAZ, TEAD4, and TEAD1 (Fig. 4c).

**Fig. 4.**
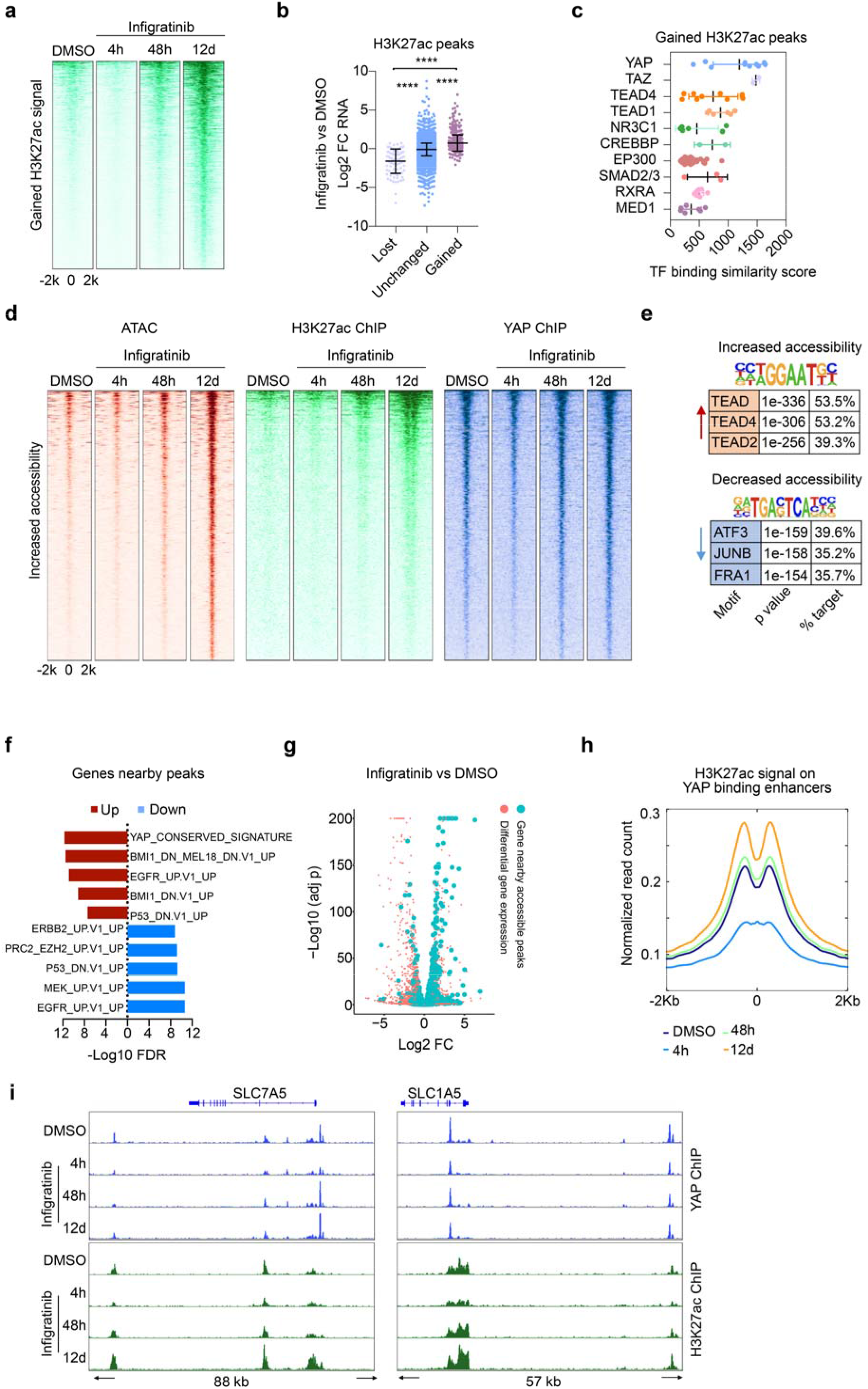
Active YAP/TEAD-dependent enhancers contribute to FGFR inhibitor resistance in TNBC. **a**, Heatmap depicting regions of significantly upregulated H3K27ac signal of CAL-120 cells upon 12d infigratinib treatment versus DMSO at indicated timepoints. **b**, RNA expression fold changes of nearest genes relative to regions of lost, unchanged, and gained H3K27ac peaks upon 12d infigratinib treatment in CAL-120 cells. Data presented as mean± SD. \*\*\*\**p*<0.0001. **c**, Plot of similarity (GIGGLE score) between published TF binding profiles and significantly increased H3K27ac ChIP-seq sites upon 12d infigratinib treatment. **d**, Heatmaps of ATAC (red), H3K27ac (green), and YAP (blue) ChIP-seq signals at significently increased ATAC-seq peaks (FDR<0.01, LFC>1.5, n=3002) upon 12d infigratinib treatment. The ChIP signals are presented in CAL-120 cells treated with DMSO and 4h, 48h, and 12d of infigratinib. **e**, Homer motif analysis of ATAC-seq peaks differentially regulated by 12d infigratinib treatment. The top 3 transcriptional factor (TF) binding motifs regulated by infigratinib are ranked by p value. **f**, Oncogenic signature gene sets enriched in the genes nearby accessible chromatin regions significantly upregulated (red) or downregulated (blue) by 12d infigratinib treatment versus DMSO. **g**, Volcano plot of –log10 adjusted p value versus log2 fold change of mRNA expression of genes differentially regulated by 12d infigratinib treatment. Genes nearby increased accessible chromatin regions (12d infigratinib vs DMSO) are colored in blue. **h**, H3K27ac signals at all YAP bound enhancers in CAL-120 cells upon DMSO and 4h, 48h, or 12d infigratinib treatment. **i**, YAP (blue) and H3K27ac (green) ChIP-seq tracks of amino acid transporters upstream of mTORC1 (SLC7A5 and SLC1A5) in CAL120 cells in the presence of DMSO or infigratinib at indicated time points.

The changes in the enhancer landscape upon FGFR inhibition led us to hypothesize that YAP/TEAD binding occurs in chromatin regulatory elements that promote transcription of target genes associated with adaptive resistance. We therefore profiled the accessible chromatin by ATAC-seq (assay for transposase-accessible chromatin using sequencing) and the YAP/TEAD bound regions by YAP ChIP-seq in CAL-120 cells upon FGFR inhibition (Supplementary Table 5). We analyzed regions of differential chromatin accessibility after 12 days of drug treatment versus control (DMSO). Among 3002 regions of significantly increased chromatin accessibility with FGFR inhibition, we also observed progressively higher H3K27ac signal at the 48 hour and 12-day time points. Overall, YAP binding events were observed in regions of increased chromatin accessibility with infigratinib treatment (Fig. 4d). Acute FGFR inhibition with 4 hour drug treatment led to decreased YAP binding, which was regained with prolonged treatment. In contrast, we observed overall decrease in the H3K27ac and YAP ChIP-seq signals in regions of decreased chromatin accessibility upon infigratinib treatment (Extended Data Fig. 4b). The majority of gained accessible regions were located at intergenic and intronic regions of the genome, with only 2.4% of gained peaks occurring within promoter regions (Extended Data Fig. 4c), suggesting that the effects we observed were primarily at enhancers.

We then utilized the unbiased motif algorithm HOMER ^31^ to identify transcription factor (TF) binding motifs enriched in regions of infigratinib-induced differentially accessible chromatin. Consistent with the H3K27ac ChIP-seq results, we found that TEAD binding motifs were significantly present in about 50% of regions of increased chromatin accessibility upon long-term FGFR inhibition (Fig. 4e). In contrast, AP-1 family binding motifs were enriched in the chromatin regions with decreased accessibility, suggesting that FGFR-MAPK downstream transcriptional events were continuously blocked by infigratinib ^32^. In line with this observation, genes nearby regions of infigratinib-increased open chromatin showed a significant enrichment of the YAP signature, whereas genes nearby decreased peaks were enriched for MEK and EGFR gene sets (Fig. 4f). By comparing open chromatin regions and the differential gene expression pattern, we found that genes nearby gained ATAC peaks were highly overlapped with upregulated RNAs upon infigratinib treatment (Fig. 4g), reflecting that the genes nearest regions of open chromatin were largely the same genes that were upregulated by FGFR inhibition.

Consistent with a central role for YAP/TEAD target genes in the feedback response to FGFR inhibition, the ChIP-seq data demonstrated that YAP binding events were also overlapped with the H3K27ac peaks after 12d of infigratinib treatment (Extended Data Fig. 4d). We next observed evidence of enhancer modulation over time at YAP bound enhancers in response to infigratinib treatment. While the H3K27ac signal was lost with 4h of treatment at these locations, it had returned to baseline by 48h and had overshot the baseline by 12 days of treatment (Fig. 4h). Taken together, these data demonstrate that FGFR inhibition acutely reduces activation of enhancers, but that the set of enhancers bound by YAP/TEAD are rapidly reactivated by 48h of FGFR inhibition, eventually exceeding their basal level of activity leading to increased expression of YAP/TEAD target genes. These results support a central role for YAP/TEAD transcriptional regulation in the response of TNBC cells to FGFR inhibition.

We then asked whether enhancer activation and YAP binding were associated with the expression of the amino acid transporters that drive increased mTORC1 signaling and resistance to infigratinib. These observations were confirmed individually for selected amino acid transporters SLC7A5 and SLC1A5, as well as canonical YAP target gene FSTL1 (Fig. 4i and Extended Data Fig. 4e). Together, these data link the enhancer activation and YAP/TEAD chromatin binding pattern to the expression of amino acid transporters that facilitate adaptive resistance to FGFR inhibition via activation of mTORC1 signaling.

### Loss of BRG1 chromatin recruitment leads to reactivation of YAP/TEAD-dependent enhancers

As YAP protein levels were not changed upon infigratinib treatment (Extended Data Fig. 5a) while the accessible enhancers were substantially altered upon long-term FGFR inhibition (Fig. 4d), we next explored whether regulators of chromatin remodeling are potentially involved in the selection of YAP/TEAD bound enhancers. By comparing the gained ATAC-seq peaks upon FGFR inhibition with the publicly available data, we found that in addition to YAP/TEAD binding sites, the binding profiles of several SWI/SNF chromatin-remodeling subunits are highly similar to the gained ATAC-seq peaks (Extended Data Fig. 5b). As ARID1A and BRG1 were among the top positively selected genes in our CRISPR KO screen upon FGFR inhibition (Fig. 1d), we hypothesize that the SWI/SNF complex might be involved in the treatment-induced chromatin remodeling.

**Fig. 5.**
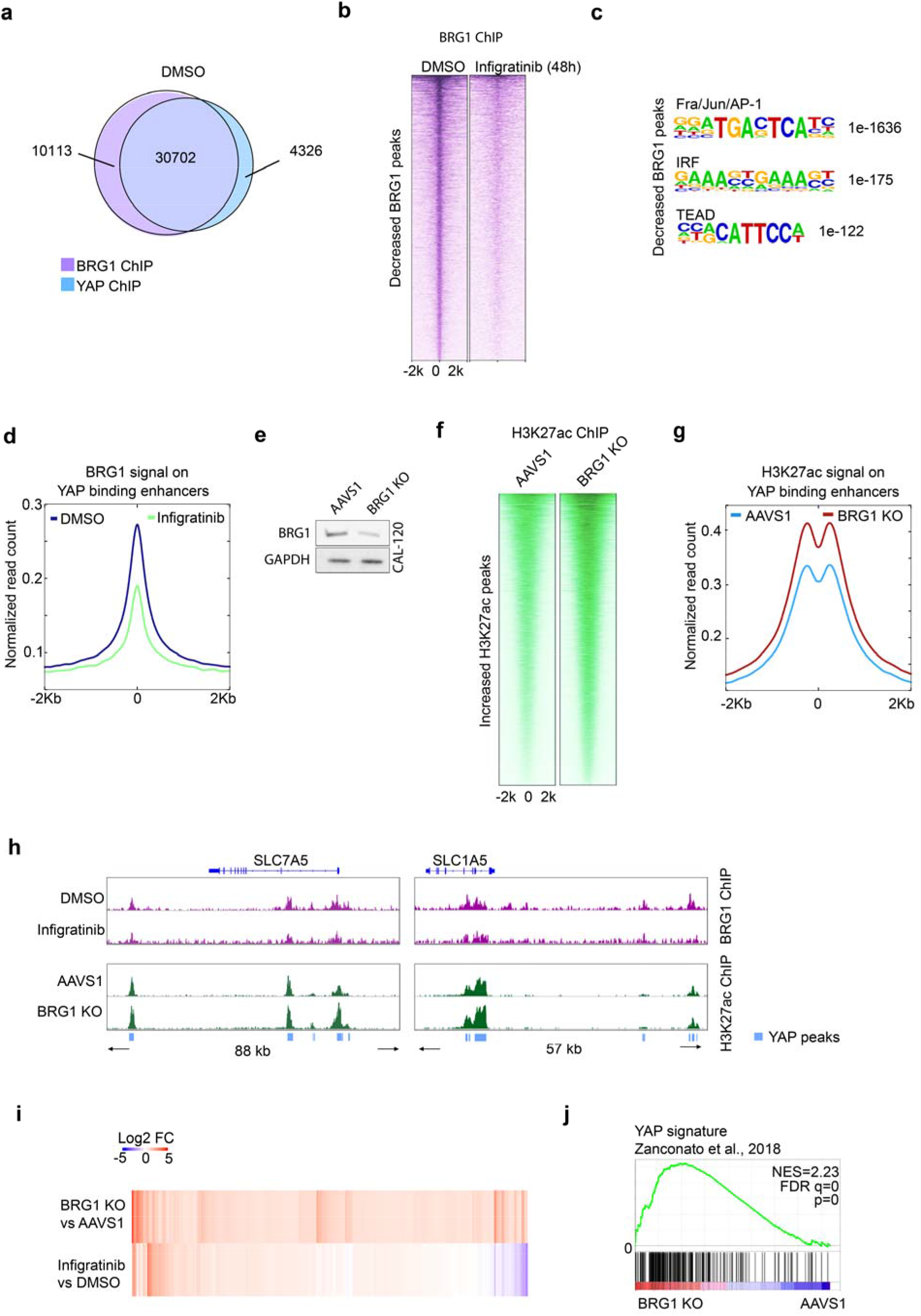
Loss of BRG1 derepresses YAP-meditated enhancer function and promotes YAP-dependent transcription. **a**, Venn diagram of overlapping BRG1 (purple) and YAP (blue) binding sites in DMSO treated CAL-120 cells. **b**, Heatmap of significantly decreased BRG1 ChIP-seq signals (n=8445) upon 48h infigratinib treatment versus DMSO. **c**, Homer motif analysis of the top 3 transcriptional factor (TF) binding motifs enriched for decreased BRG1 peaks in b. **d**, BRG1 ChIP-seq signals at all YAP bound enhancers defined in figure 4h in CAL-120 cells upon DMSO or 48h infigratinib treatment. **e**, Immunoblot of BRG1 and GAPDH in CAL-120 cells knocked out for AAVS1 control or BRG1. **f**, Heatmap of significantly increased H3K27ac peaks (n=14620) in CAL-120 cells depleted for BRG1 versus AAVS1 control. **g**, H3K27ac ChIP-seq signals at all YAP bound enhancers defined in figure 4h in CAL-120 cells depleted for AAVS1 or BRG1. **h**, Upper, BRG1 (purple) ChIP-seq tracks of SLC7A5 and SLC1A5 in CAL120 cells in the presence of DMSO or 48h infigratinib. Lower, H3K27ac (green) ChIP-seq tracks of SLC7A5 and SLC1A5 in AAVS1 and BRG1 depleted cells. YAP DNA binding regions are marked in blue. **i**, Heatmap representing Log2 fold change of the mRNA levels of the 1211 significantly upregulated genes in BRG1 knockout versus AAVS1 knockout CAL-120 cells and the Log2 fold change of these same genes in CAL-120 cells treated with infigratinb for 48h versus DMSO control. **j**, GSEA analysis reveals the YAP target gene signature is significantly enriched in CAL120 cells depleted of BRG1.

To explore whether the feedback activation of enhancers is determined by interplay between YAP/TEAD and the BRG1-containing SWI/SNF complex, we performed ChIP-seq for BRG1 in CAL-120 cells in presence or absence of infigratinib at 48h when the feedback response was first evident. The BRG1 and YAP1 cistromes substantially overlapped in untreated (DMSO) CAL-120 cells (Fig. 5a). However, treatment with infigratinib resulted in a dramatic alteration of BRG1 binding (Extended Data Fig. 5c). The majority of differentially BRG1 bound regions were decreased upon FGFR inhibition (Fig. 5b). Infigratinib-suppressed BRG1 binding sites are highly enriched for Fra/Jun/AP-1 (39.3%), TEAD (26.2%) and IRF (6.9%) motifs (Fig. 5c), indicating many YAP binding sites lose the BRG1 binding activity upon FGFR inhibition. BRG1 binding at YAP bound enhancers was markedly inhibited by drug treatment (Fig. 5d). This finding suggests that BRG1 is recruited to YAP/TEAD binding enhancers at normal chromatin state and abrogation of BRG1 function is associated with adaptive resistance to infigratinib.

Previous work has shown that the SWI/SNF complex can interact with YAP/TAZ and suppress YAP mediated transcription ^33^. We hypothesized that loss of BRG1 activity might promote the activation of YAP/TEAD dependent enhancers. BRG1 was knocked out in CAL-120 cells using CRISPR-mediated gene editing (Fig. 5e). We performed ChIP-seq of H3K27ac in control (AAVS1) and BRG1 knockout (BRG1 KO) cells. The H3K27ac signal was increased globally in the BRG1 KO cells, supporting the hypothesis that loss of SWI/SNF is involved in enhancer reactivation (Fig. 5f and Extended Data Fig. 5d). Notably, the genes nearby infigratinib-reduced BRG1 binding sites are highly overlapped with genes close to gained H3K27ac signal in BRG1 KO cells (Extended Data Fig. 5e). Furthermore, the H3K27ac binding signal on YAP/TEAD enriched enhancers was increased in the BRG1 KO cells (Fig. 5g). In line with this, infigratinib treatment repressed BRG1 binding at the enhancer regions of SLC7A5 and SLC1A5 that overlap with YAP-bound sites, whereas H3K27ac signal was increased at the enhancer regions of SLC7A5 and SLC1A5 upon BRG1 knockout (Fig. 5h). These data confirm that loss of BRG1 leads to reactivation of YAP/TEAD bound enhancers.

To identify whether BRG1-mediated enhancer activation can result in a similar transcriptional pattern as FGFR inhibition, we performed RNA-seq analysis in control and BRG1 KO CAL-120 cells. We observed 1211 genes were significantly induced by BRG1 depletion, and majority of these genes were also consistently increased by infigratinib treatment (Fig. 5i). The gene expression profile of BRG1 KO cells was significantly enriched in YAP target genes ^24^ (Fig. 5j), suggesting BRG1 antagonizes YAP transcriptional regulation in TNBC cells. Overall, we found FGR inhibition represses chromatin recruitment of BRG1 leading to the remodeling of regulatory regions bound by YAP/TEAD causing an epigenetic switch to YAP transcriptional dependency.

## Discussion

Available data from FGFR inhibitor clinical trials suggest overall response rates in FGFR-aberrant solid tumors of approximately 25-40% ^11, 19, 34^, suggesting that the identification of resistance mechanisms to FGFR inhibition is critical to maximizing the number of patients that could benefit from this targeted therapy. In this study, we identified mTOR and YAP loss as sensitizers of FGFR inhibition, while ARID1A or BRG1 depletion can engender drug resistance. We demonstrated that adaptive resistance to FGFR inhibition occurs via inhibition of SWI/SNF-regulated epigenetic state, resulting in reactivation of YAP dependent enhancers that in turn promote the amino acid transport that is sensed by the mTORC1 pathway leading to cell growth. This FGFR inhibitor activated feedback loop that limits the efficacy of these inhibitors can be reversed by mTORC1 inhibition (Fig. 6).

**Fig. 6.**
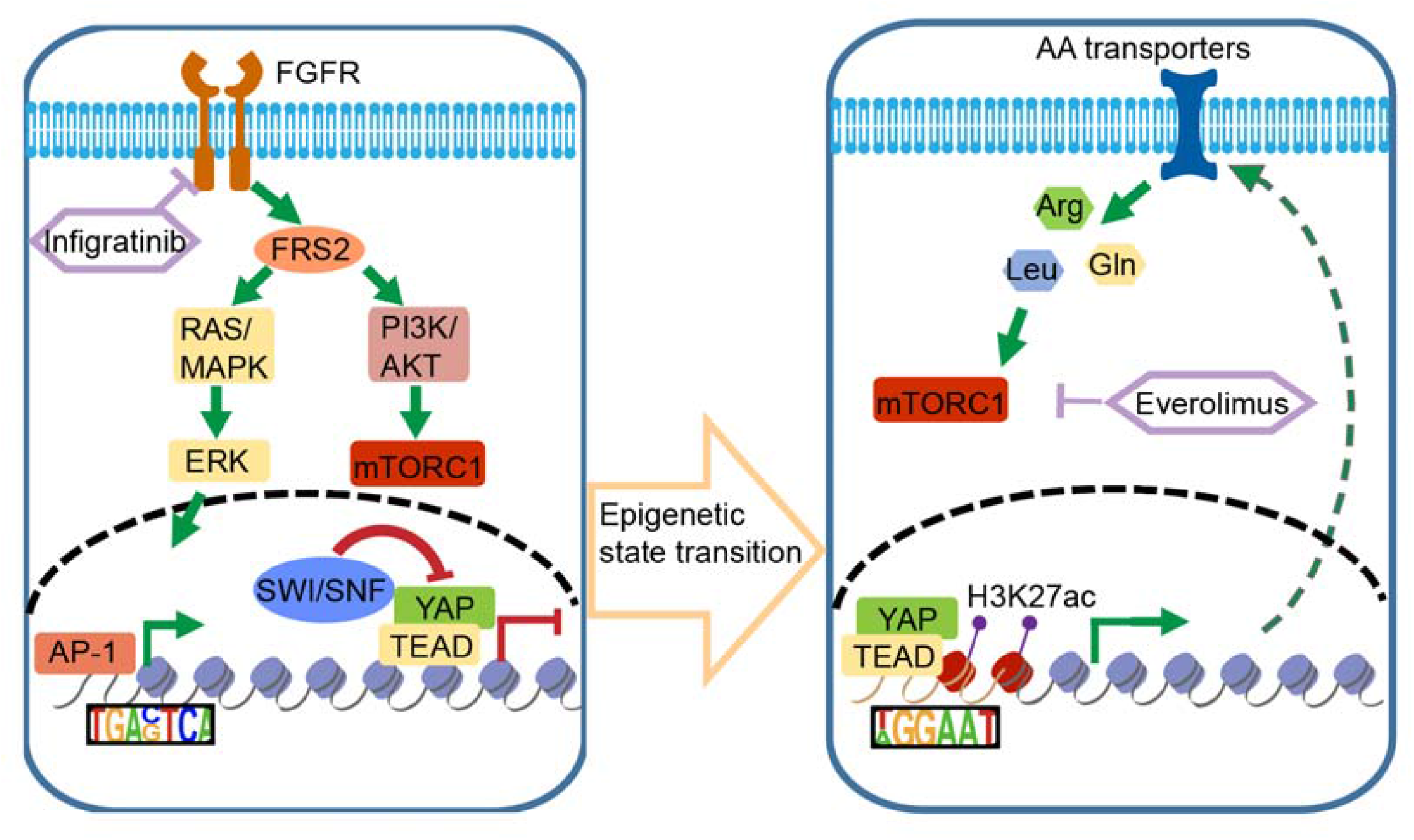
Epigenetic feedback loop driving adaptive resistance to FGFR inhibition.

Metabolic reprogramming is one hallmark of cancer progression, potentially uncovering new vulnerabilities. The solute carrier (SLC) transporters serve as the plasma membrane sensors controlling mTORC1 activity through regulation of intracellular levels of amino acids. However, these transporters are rarely mutated in human cancers, indicating the intracellular level of amino acids is regulated through changes in their expression ^35^. In addition, therapies targeting transporters have yet to be investigated in clinical trials. We found that FGFR inhibition upregulated the expression of a set of amino acid transporters, implicating the important role of amino acid dependent mTORC1 signaling as a novel mechanism of therapeutic resistance. In this study, the combined withdrawal of amino acids was, at least in part, able to suppress the feedback loop leading to mTORC1 activation. Although mTOR feedback activation has been observed in other cancer drug resistant settings, these have commonly been secondary to autocrine growth factor receptor activation rather than amino acid sensing ^36^. Our findings thus reveal a mechanism of crosstalk between Hippo and mTOR signaling and suggest an additional potential therapeutic target upstream of mTOR.

Several prior reports have identified altered enhancer landscape as a mechanism of adaptive resistance to targeted therapies ^37, 38^. We observed remodeling of accessible enhancers upon FGFR inhibition in TNBC cells. This epigenetic shift is governed by loss chromatin binding at YAP-bound enhancers of BRG1, a core component of a SWI/SNF remodeling complex. Loss of BRG1 binding leads to increased levels of H3K27 acetylation and activation of these YAP-bound enhancers and the expression of a YAP-dependent transcriptional program. Our finding is consistent with a prior report showing that YAP and its downstream effectors are highly associated with aggressive cellular phenotypes including cancer stem cell properties and metastasis in breast cancer cells ^22^. Our findings also align with reports of physical interaction between ARID1A and YAP ^33^. The interplay between SWI/SNF and YAP transcriptional regulation may thus play an important role in tumor progression and drug response in basal-like triple negative breast cancer.

Overall, our findings reveal a feedback mechanism underlying the limited clinical efficacy of FGFR inhibitors in TNBC that based on the presence of FGFR alterations would otherwise be predicted to be FGFR dependent. This mechanism involves the loss of the BRG1 chromatin remodeling factor from a set of YAP dependent enhancers leading to reactivation of a YAP transcriptional program. This program includes upregulation of amino acid transporters leading to increased amino acid sensing by mTORC1.

## Acknowledgments

This work was supported by the Ludwig Center at Harvard Medical School to M.B and A.T. We thank members from Brown lab and Liu lab for helpful discussions and technical help. We thank the Molecular Biology Core Facilities (MBCF) at Dana-Farber Cancer institute (DFCI) for help with next generation sequencing. We thank Steven Gygi and Rachel Rodrigues for multiplexed proteomic analysis.

## Author contributions

Y.L. and M.B. designed experiment; Y.L., X.Q., X.W., H.L., R.C.G., T.X. A.F., K.L., P.K., A.J., R.V., P.C., K.L.J., M.M. performed methodology, investigation, and validation, Y.L., X.Q., R.C.G., and Y.X; performed software and formal analysis, K.C., P.C., Q.N., D.W., H.W.L, X.S.L, A.T., and M.B. provided resources and supervision, Y.L., X.Q., A.K.T., A.T., and M.B. wrote the manuscript. All authors read and corrected the manuscript.

