## supplemental figures for "FGFR inhibitor mediated dismissal of SWI/SNF complexes from YAP-dependent enhancers induces adaptive therapeutic resistance"

Fig S1

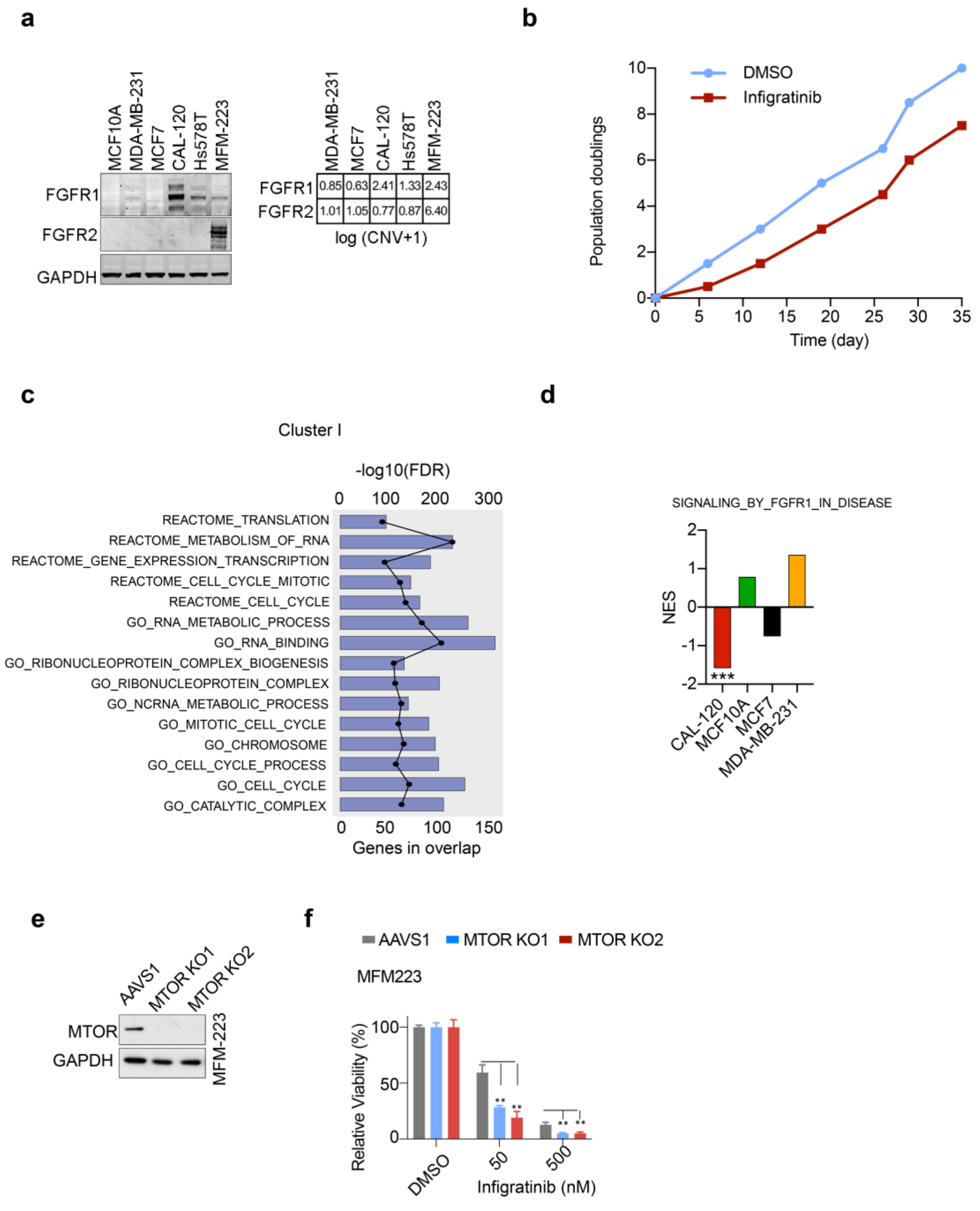


**Extended Data Fig. 1: A genome-wide CRISPR knock out screen identifies genetic dependencies in FGFR-aberrant TNBC.**

**a,** Left, immunoblot of FGFR1 and FGFR2 protein levels in MCF10A, MDA-MB-231, MCF7, CAL-120, Hs578T and MFM223 cells. Right, copy number variation (CNV) of indicated cell lines from Cancer Cell Line Encyclopedia (CCLE) dataset. **b,** Population doublings of CAL-120 cells treated with DMSO or 300nM infigratinib for 35 days during genome-wide CRISPR screen. **c,** Gene ontology analysis for cluster 1 essential genes (essential in multiple cell lines) from Figure 1D. Each of the top enriched gene sets are from the GO and reactome gene sets in the Molecular Signature Database (MSigDB). **d,** GSEA normalized enrichment score (NES) of FGF signature for essential genes in the indicated cell line, asterisk indicates statistically significant enrichment (****p*<0.001). **e,** Immunoblot of mTOR protein levels in MFM-223 cells after introduction of CRISPR guide RNAs targeting AAVS1 (control) or mTOR. **f,** The relative cell viability of MFM-223 cells depleted of mTOR versus AAVS1 (control) in the presence of DMSO, 50 nM, or 500 nM infigratinib for 6 days.

Fig S2

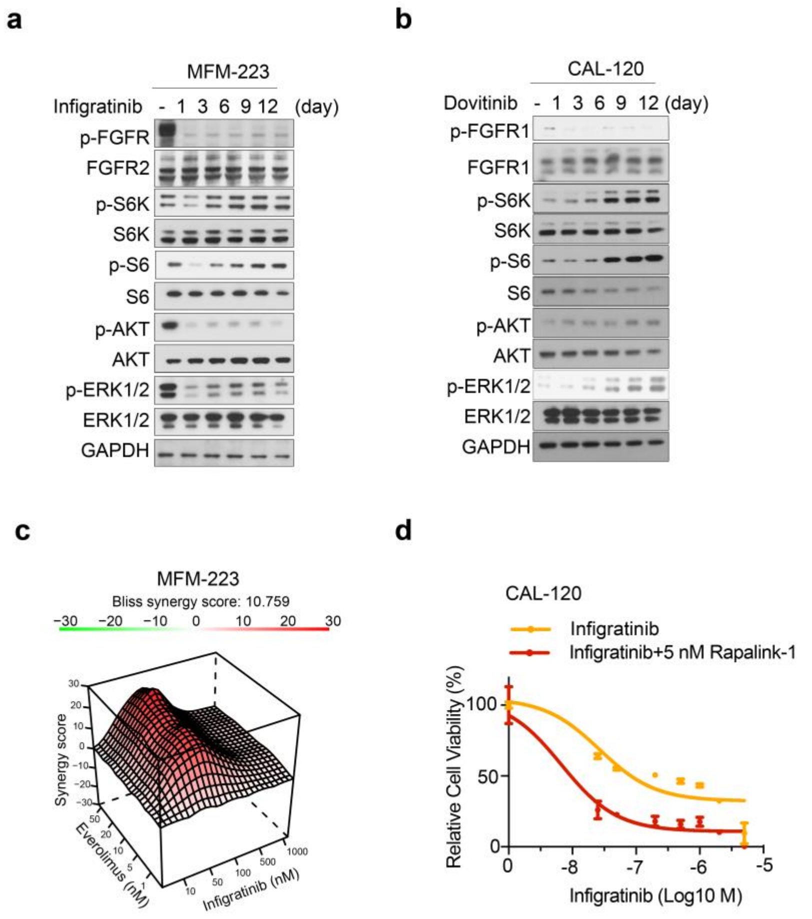


**Extended Data Fig. 2: Role of mTORC1 signaling in adaptive resistance to FGFR inhibitors.**

**a,** Immunoblot of FGFR, ERK1/2, AKT, S6K, and S6 phosphoprotein and total protein levels in MFM-223 cells treated with 50 nM infigratinib at the indicated timepoints. **b,** Immunoblot of FGFR1, ERK1/2, AKT, S6K, and S6 phosphoprotein and total protein levels in CAL-120 cells treated with 300 nM dovitinib at the indicated time points. **c,** Bliss synergy model of combination infigratinib and everolimus treatment in MFM-223 cells. **d,** Relative cell viability of CAL-120 cells treated with infigratinib in a dose-dependent manner alone or in combination with 5 nM rapalink-1 for 6 days.

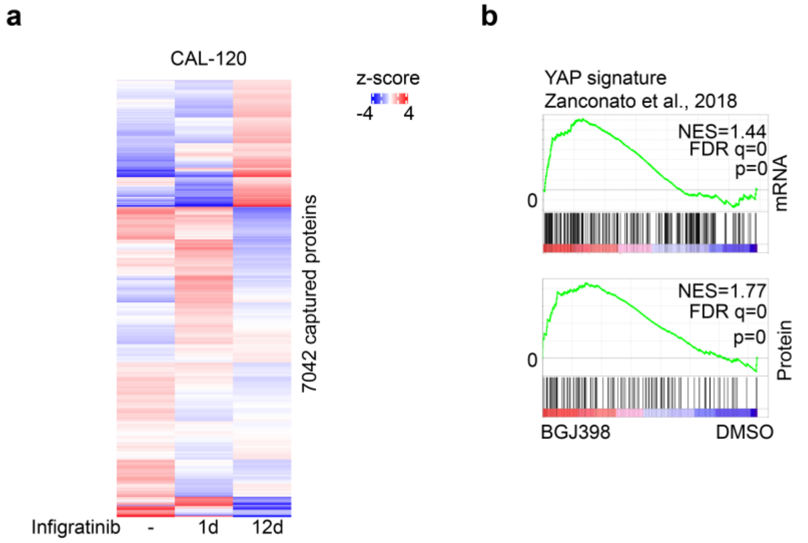


**Extended Data Fig.3: Increased expression of YAP target genes and amino acid transporters are driven by FGFR** inhibition. **a,** Heatmap of protein expression levels detected by multiplexed proteomics in CAL-120 cells upon DMSO, 1d, or 12d of 300 nM infigratinib treatment. Rows are clustered by k-means analysis. **b,** GSEA analysis reveals a YAP signature is significantly enriched in genes differentially regulated by 12d of infigratinib treatment in CAL-120 cells.

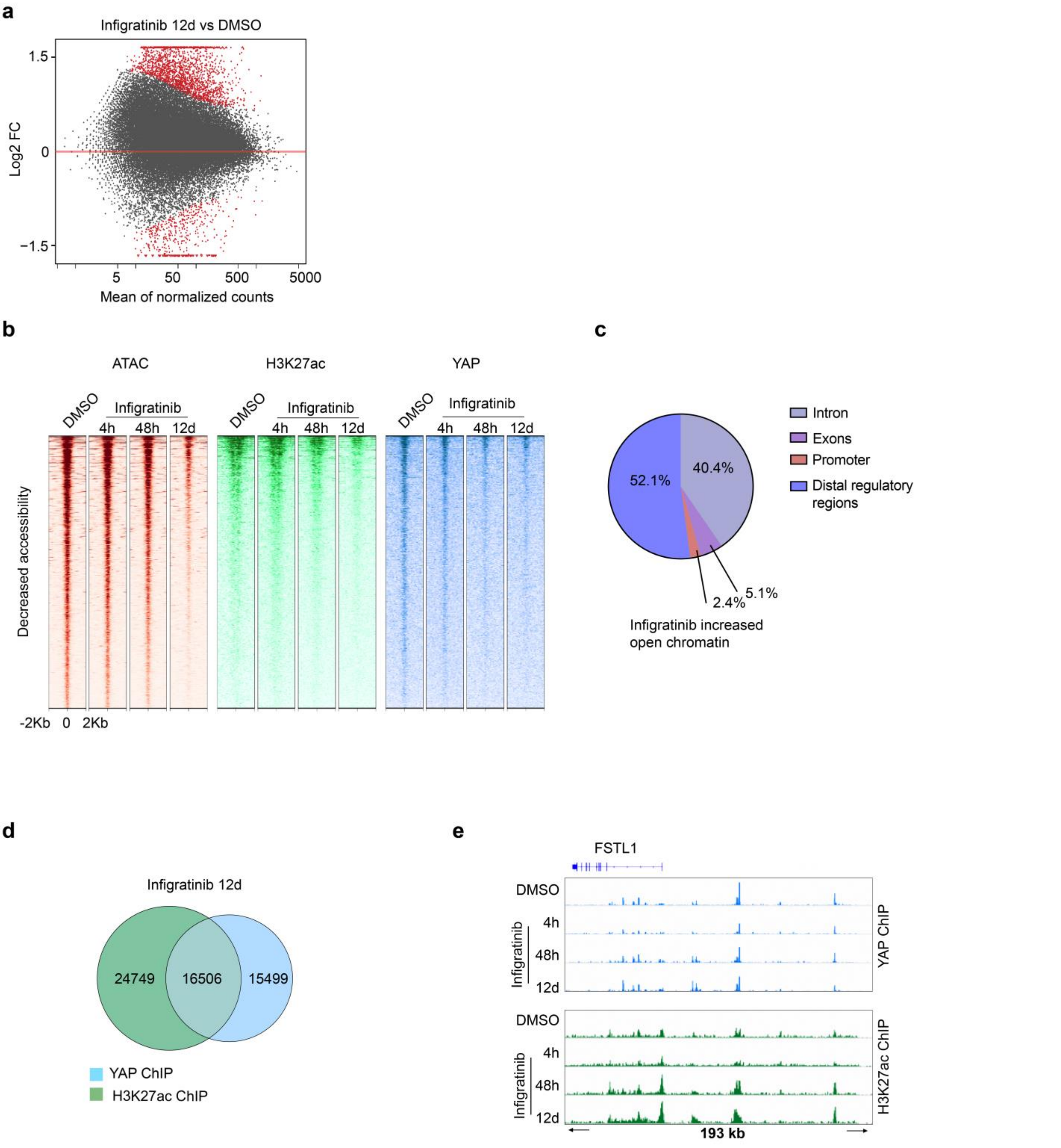


**Extended Data Fig. 4: FGFR inhibition regulates enhancer landscape in TNBC cells.**

**a,** MA plot of differentially regulated H3K27ac ChIP-seq peaks in CAL-120 cells treated with 12d infigratinib versus DMSO. Mean of normalized counts of each peaks are plotted on the x-axis and log2 fold changes of differential peaks are plotted on the y-axis. Significantly changed peaks (FDR<0.05) are marked in red. **b,** Heatmaps of ATAC-seq, H3K27ac and YAP ChIP-seq signals at regions of decreased chromatin accessibility (FDR<0.01, LFC<-1.5, n=2798) upon 12d infigratinib treatment. **c,** Genomic distribution of regions of increased chromatin accessibility upon 12d infigratinib treatment. **d,** Venn diagram depicting overlap between H3K27ac peaks (green) and YAP (blue) DNA binding peaks after 12d infigratinib treatment. **e,** YAP (blue) and H3k27ac (green) ChIP-seq tracks around YAP target gene FSTL1 in CAL120 cells in the presence of DMSO or infigratinib at indicated time points.

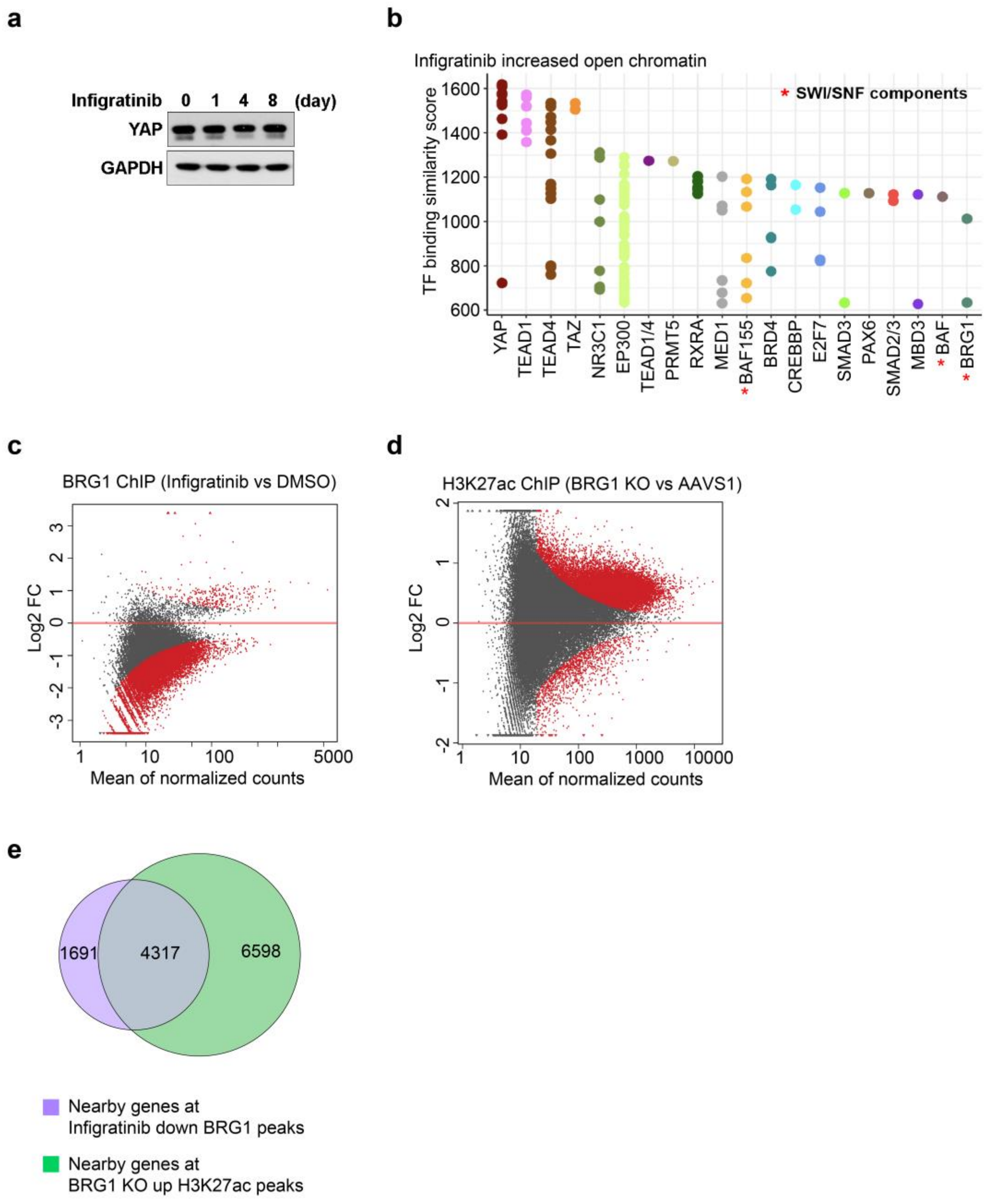


**Extended Data Fig. 5: Infigratinib treatment decreases BRG1 chromatin binding and mediates enhancer reactivation.**

**a,** Immunoblot of YAP protein levels in CAL-120 cells treated with 300 nM infigratinib at the indicated timepoints. **b,** Plot of similarity (GIGGLE score) between published binding profiles of SWI/SNF complex components and regions of increased chromatin accessibility in CAL-120 cells upon 12d infigratinib treatment. SWI/SNF subunits are marked by *. **c,** MA plot of differential BRG1 ChIP-seq signals in CAL-120 cells treated with 48h infigratinib versus DMSO. Mean of normalized counts of each peaks are plotted on the x-axis and log2 fold changes of peaks are plotted on the y-axis. Significantly changed peaks (FDR<0.05) are marked in red. **d,** MA plot of differential H3K27ac ChIP-seq signals in CAL-120 cells depleted of BRG1 relative to AAVS1 control. Significantly changed peaks (FDR<0.05) are marked in red. **e,** Venn diagram illustrating the overlap in the nearby gene to lost BRG1 binding sites upon 48h FGFR inhibition from figure 5b and nearby gene to gained H3K27ac signal after BRG1 depletion from figure 5f.
